# Structure of the Ebola virus nucleoprotein – RNA complex

**DOI:** 10.1101/548313

**Authors:** Robert N. Kirchdoerfer, Erica Ollmann Saphire, Andrew B. Ward

## Abstract

Ebola virus is an emerging virus capable of causing a deadly disease in humans. Replication, transcription and packaging of the viral genome is carried out by the viral nucleocapsid. The nucleocapsid is a complex of the viral nucleoprotein, RNA and several other viral proteins. The nucleoprotein NP forms large, RNA-bound, helical filaments and acts as a scaffold for additional viral proteins. The 3.1 Å single-particle cryo-electron microscopy structure of the nucleoprotein-RNA helical filament presented here resembles previous structures determined at lower resolution while providing improved molecular details of protein-protein and protein-RNA interactions. The higher resolution of the structure presented here will facilitate the design and characterization of novel and specific Ebola virus therapeutics targeting the nucleocapsid.

**Synopsis:** The 3.1 Å single-particle cryo-electron microscopy structure of the RNA-bound, Ebola virus nucleoprotein helical filament provides molecular details of protein-protein and protein-RNA interactions.

## 1. Introduction

Ebola virus is capable of causing a disease manifesting a severe hemorrhagic fever with case fatality rates as high as 90% (*Ebola haemorrhagic fever in Zaire, 1976*, 1978). Though generally outbreaks are geographically isolated, the outbreak of 2014-2015 was responsible for over 28,000 cases and spread between countries (Hersey *et al.*, 2015). As a nonsegmented negative-sense RNA virus, the RNA genome of Ebola virus is encapsidated by a viral nucleoprotein throughout the virus life cycle. This protein-RNA complex serves as a scaffold for additional viral proteins including the viral polymerase to carry out viral transcription and replication. The complex of RNA, nucleoprotein and these additional viral proteins forms the nucleocapsid and is a structural component of Ebola virions. Full length Ebola virus nucleoprotein is 739 amino acids in length. However, the N-terminal 450 amino acids (NP 1-450) are necessary and sufficient for protein oligomerization and RNA-binding (Watanabe *et al.*, 2006) and this N-terminal region of the protein forms helical filaments resembling NP in virions (Bharat *et al.*, 2012).

The determinants of NP oligomerization and RNA binding have been the focus of several structural studies. Crystal structures and hydrogen/deuterium exchange mass spectrometry have shown that the N-terminal region of NP possesses a folded core flanked by disordered regions (Kirchdoerfer *et al.*, 2015, Leung *et al.*, 2015). The folded core of NP possesses N- and C-terminal lobes and is primarily alpha-helical in structure. The available crystal structures of Ebola NP were all determined using monomeric NP (Dong *et al.*, 2015, Kirchdoerfer *et al.*, 2015, Leung *et al.*, 2015). For two of these structures, NP is bound to a peptide of viral protein 35 (VP35) (Kirchdoerfer *et al.*, 2015, Leung *et al.*, 2015). The binding of this VP35 peptide is thought to be important for maintaining a monomeric state of NP, and preventing the premature and non-specific encapsidation of host RNA (Kirchdoerfer *et al.*, 2015, Leung *et al.*, 2015, Leyrat *et al.*, 2011).

After the VP35 peptide has been removed, NP oligomerizes into a linear polymer capable of encapsidating RNA. Oligomerization of NP is mediated by protein regions N- and C-terminal to the two-lobed folded NP 1-450 core (Kirchdoerfer *et al.*, 2015, Leung *et al.*, 2015). This oligomerization of NP causes a conformational change between the two lobes creating a high-affinity binding site for RNA (Kirchdoerfer *et al.*, 2015, Sugita *et al.*, 2018). Cryo-electron tomography studies of Ebola virus nucleocapsids as well as Ebola virus NP 1-450 demonstrated the left-handed helical nature of the NP filaments, as well as the first structural evidence of NP oligomerization regions mediating contacts between NP monomers (Wan *et al.*, 2017). The limited resolution of this earlier cryoET work, however, precluded clear elucidation of molecular interactions among NP and with RNA. A more recent study using single particle cryo-electron microscopy of the NP 1-450 determined the structure of this helical filament at 3.6 Å resolution (Sugita *et al.*, 2018). This structure revealed molecular interactions as well as conformational changes between the monomeric, chaperoned NP and the oligomeric, RNA-bound NP. Here, we have used single-particle cryoEM to determine the structure of the NP 1-450 helix at an improved resolution of 3.1 Å.

## 2. Materials and Methods

### 2.1. Expression of NP helical filaments

NP filaments were prepared similar to previously described methods (Bharat *et al.*, 2012). 0.5 mg NP 1-450-pDisplay plasmid without any additional peptide tags was diluted into 20 mL of OPTI-MEM and then filtered through a 0.22 µm filter. To this solution was added 1.5 mg of polyethylenimine in 5 mL of OPTI-MEM. The DNA-PEI complexes were allowed to incubate for 20 minutes at room temperature and then added to 1 L of 293F cells at 1.1×10^6^ cells/mL growing in 293Freestyle media in a 2 L baffled flask. Transfected cells were incubated at 37°C with 8% CO_2_ at 130 rpm for 48 hours.

### 2.2. Purification of NP helical filaments

Cells were collected by centrifugation at 1,000 × g for 15 minutes and then resuspended in 8 mL 25 mM TrisCl, 150 mM sodium chloride, 1 mM calcium chloride and 5 mM beta-mercaptoethanol. The cells were then lysed by the addition of 10 mL of 25 mM TrisCl pH 7.4, 150 mM sodium chloride, 1 mM calcium chloride, 0.2% Igepal CA-630 and 5 mM beta-mercaptoethanol. The lysate was clarified by centrifugation at 3,200 × g for 20 minutes.

The NP helical filaments were isolated by loading the cleared lysate onto a double cushion of 5 mL of 20% and 5 mL of 90% sucrose. The sucrose cushions were spun in an ultracentrifuge using a Beckman SW28 rotor at 28,000 rpm for three hours. The sucrose cushions were fractioned by bottom puncture and ∼1 mL manually collected fractions were analyzed by SDS-PAGE. Fractions containing NP 1-450 were dialyzed into 500 mL of 25 mM TrisCl, 150 mM sodium chloride, 1 mM calcium chloride and 5 mM beta-mercaptoethanol using a 1 MDa molecular weight cutoff dialysis tubing overnight with one buffer change to remove sucrose. 3 mL of the dialyzed protein was then loaded onto one of two discontinuous cesium chloride gradients containing 24 mL 25% and 10 mL 60% w/v. The cesium chloride gradients were spun in a SW28 rotor at 28,000 rpm for four hours. Gradients were then manually fractionated after bottom tube puncture and fractions were analyzed with SDS-PAGE before overnight dialysis of fractions containing NP 1-450 to remove cesium chloride.

NP 1-450 was concentrated by pelleting the dialyzed protein (∼ 3 mL) through 400 µL of a 20% sucrose cushion using a Beckman TLA100.3 rotor at 48,000 rpm for 1 hour. The pellet was resuspended in 400 µL 25 mM TrisCl pH 7.4, 150 mM sodium chloride, 1 mM calcium chloride and 5 mM beta-mercaptoethanol. Measuring the UV absorbance of the resuspended sample yielded an A_280_ of 3.8 with an A_260_/A_80_ ratio of 1.8.

### 2.3. Cryo-electron microscopy data collection and preprocessing

UltraAu Foil R 1.2/1.3 Au 300 mesh grids (Quantifoil) were plasma cleaned for 7 s with an Ar/O_2_ gas mix. 4 µL of sample was blotted onto the grids and grids were blotted for 4 s using a Vitrobot IV (Thermo Fisher) before plunge freezing into liquid ethane. Data was collected using Leginon (Suloway *et al.*, 2005) on a Talos Arctica (Herzik *et al.*, 2017) (Thermo Fisher) operating at 200 kV and a K2 Summit detector (Gatan) in counting mode using a magnification of 43,478 and at a dose rate of 5.06 e/pix/sec, collecting 200 ms frames for a 13 second exposure and a total dose of 49.7 e^-^/Å^2^. Frame alignments were performed using MotionCorr2 (UCSF) (Zheng *et al.*, 2017). CTF estimates were performed with GCTF (Zhang, 2016) and images with estimated resolutions lower than 5 Å were removed.

### 2.4. Data processing and refinement

Filaments were picked manually, and particles were extracted from filaments using a box size of 360 pixels at a pixel size of 1.15 Å/pixel and an interbox distance of 100 Å (He & Scheres, 2017). The resulting stack of particles was cleaned by 2D classification in Relion 2.1 (Kimanius *et al.*, 2016) where poorly aligned particles were excluded from subsequent processing. 3D refinement was initiated with a bead model produced using the relion_helix_toolbox (He & Scheres, 2017) and helical parameters estimated from a previous reconstructions of the Ebola virus nucleocapsid (Bharat *et al.*, 2012, Wan *et al.*, 2017). These same helical parameters were used as starting helical parameters for 3D refinement and these parameters were allowed to optimize during refinement.

Atomic coordinates were built into density accounting for the asymmetric unit of the helix using Coot (Emsley *et al.*, 2010) and a monomeric Ebola virus NP structure, 4ZTG.pdb as a starting model. The 3.6 Å model previously published was used for comparison (Sugita *et al.*, 2018), 5Z9W.pdb. The coordinates were refined against the map using RosettaRelax (DiMaio *et al.*, 2015). Final coordinates were evaluated using Molprobity (Williams *et al.*, 2018) and EMRinger (Barad *et al.*, 2015). Data collection and processing as well as model refinement statistics are presented in Table 1.

**Table 1.** Data collection and coordinate model refinement statistics.

| Data and reconstruction statistics |  |
| --- | --- |
| EMDB | EMD-0522 |
| Microscope | Talos Actica |
| Voltage (kV) | 200 |
| Detector | Gatan K2 Summit |
| Dose rate (e<sup>-</sup>/pix/sec) | 5.06 |
| Exposure (s) | 13 |
| Dose (e<sup>-</sup>/Å<sup>2</sup>) | 49.7 |
| Frames | 65 |
| Defocus Range (μm) | 0.6 – 2.0 |
| Filaments | 2,143 |
| Particles | 24,609 |
| B-factor (Å<sup>2</sup>) | -94 |
| Resolution (Å) | 3.1 |
| Coordinate model refinement |  |
| PDB | 6NUT |
| Residues |  |
| Amino acids | 379 |
| Nucleotides | 6 |
| RMSD Bonds (Å) | 0.016 |
| RMSD Angles (°) | 1.35 |
| Ramachandran |  |
| Favored (%) | 99.2 |
| Allowed (%) | 0.8 |
| Outliers (%) | 0.0 |
| Rotamer Outliers (%) | 0.3 |
| Clash score | 2.89 |
| Molprobity score | 1.08 |
| EM Ringer Score | 3.15 |

## 3. Results

The NP helical filaments have a barbed appearance in the aligned, dose-weighted micrograph movies (Fig. 1). 2D alignment of particles revealed classes with clear secondary structure features. The refined 3D particle orientations demonstrate a predominance of views perpendicular to the helical axis, as expected, given the orientations of the helical filaments picked in the raw micrographs (Fig. 2A). The refined helical parameters for the final map are −14.71° twist and a 2.84 Å rise per NP protomer. The reconstructed map has a resolution of 3.1 Å estimated using a gold-standard Fourier shell correlation cutoff of 0.143. Local resolution estimation shows a resolution range of 3.0-3.8 Å (Fig. 2), with the folded core of the NP exhibiting the highest resolution, while peripheral, surface exposed regions reconstructed to lower resolution, possibly indicating local mobility or conformational pleiomorphism.

**Figure 1:**
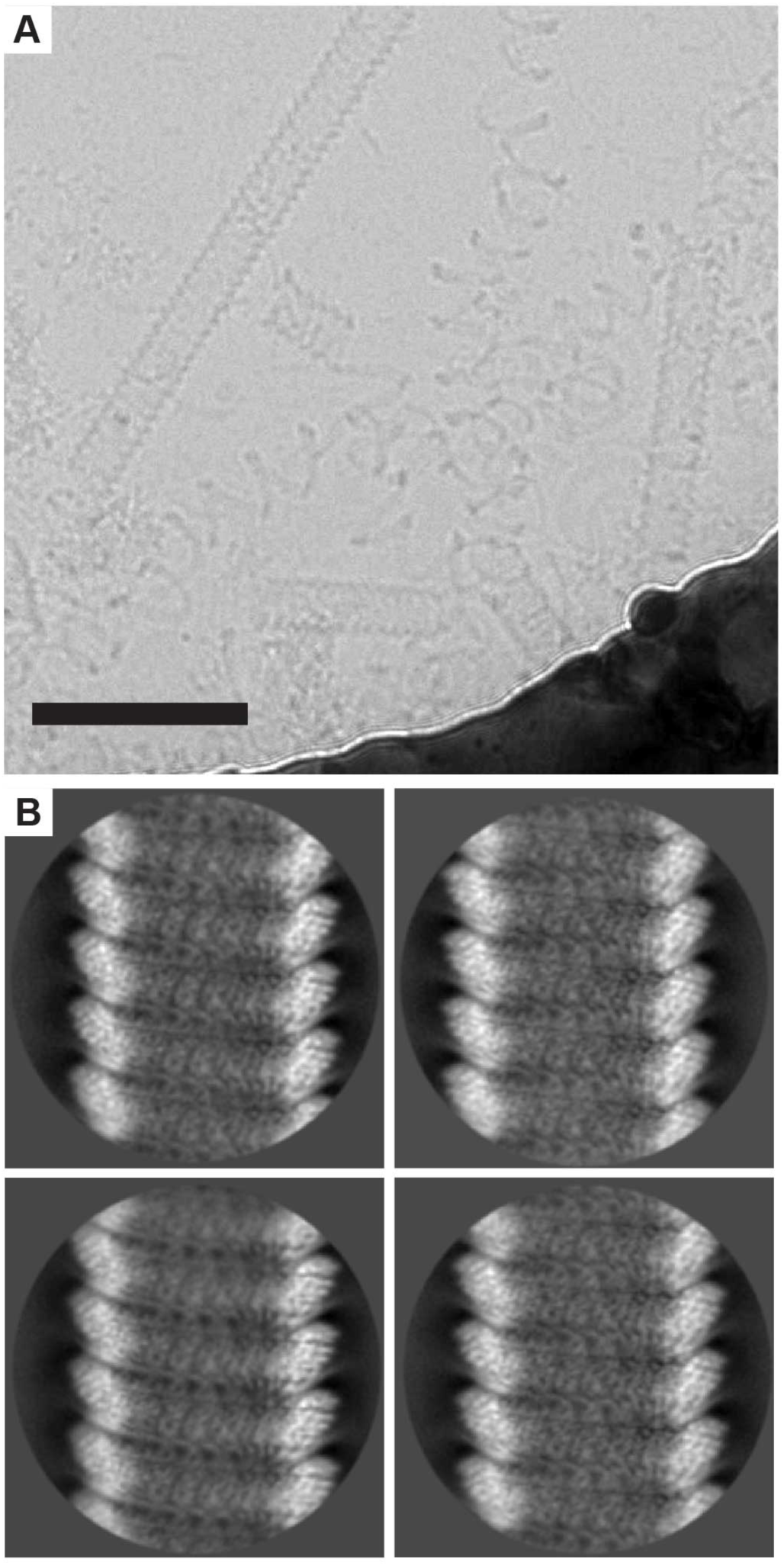
Image acquisition and processing by two-dimensional alignment.) A) Raw micrograph movies were aligned and dose weighted. Scale bar is 100 nm. B) 2D alignment of picked particles shows the helical nature of the particles with evidence of secondary structure.

**Figure 2:**
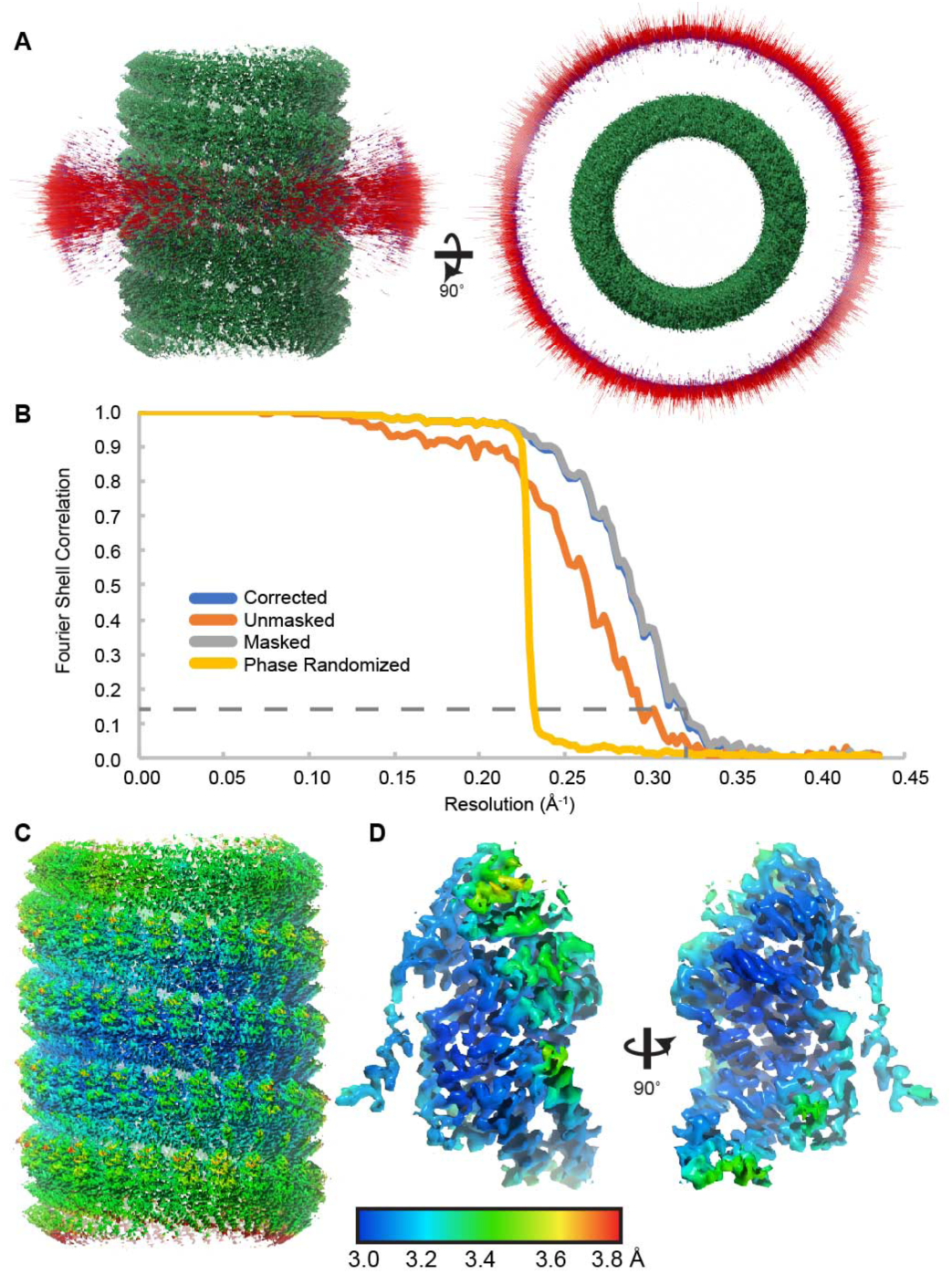
Particle angular distribution and map resolution. A) The angular distribution of refined particle orientations indicates a predominance of views (red) perpendicular to the filament axis. B) Fourier shell correlation plots indicated a resolution of 3.1 Å using a gold-standard FSC cutoff of 0.143 indicated by the dashed line. Local resolution of the NP helical filament (C) and an isolated NP protomer (D) indicate the high-resolution folded core of NP with lower-resolution peripheral regions.

The high resolution of the map enabled building and refinement of an atomic model of the Ebola virus NP bound to RNA. Side-chain density for most amino acids is clearly evident and we observe density for six nucleotides of RNA similar to previous studies (Sugita *et al.*, 2018) (Fig. 3). Because NP is believed to bind to RNA without sequence specificity, the RNA nucleotides seen in the map represent the average of bound nucleotides. We modeled this density as a polymer of six adenosine residues since, as expected, none of these residues makes sequence specific contacts.

**Figure 3:**
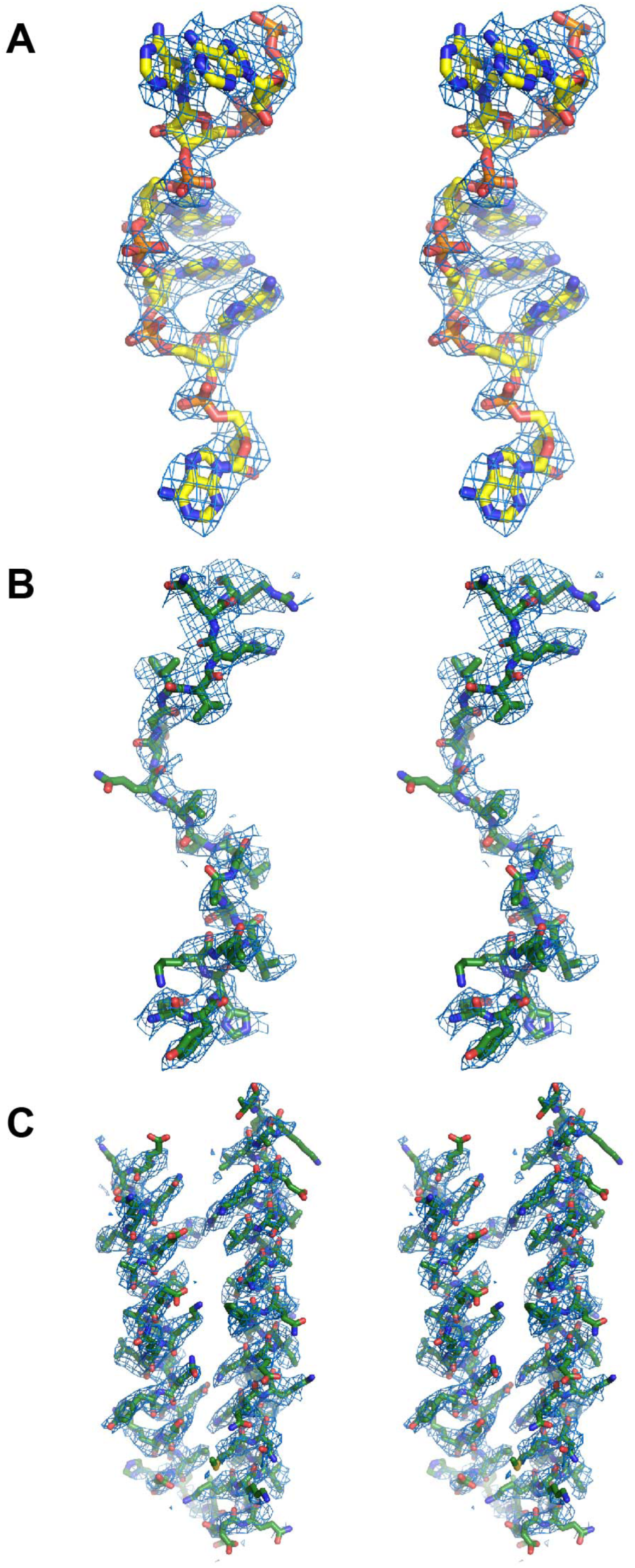
Fit of model to reconstructed density. Stereo views of the NP-RNA helical filament focusing on (A) the bound RNA, (B) the N-terminal oligomerization arm and (C) the C-terminal oligomerization helices.

The coordinate model presented here overlays well with the previously published 3.6 Å structure of NP bound to RNA (Sugita *et al.*, 2018) (Fig 4). There does appear to be a 2.2% isotropic stretching between the previously published map and the map presented here. We attribute this to issues in pixel size calibration of the previously collected data. However, this effect is small and unlikely to impact conclusions drawn from this previous study. Future antiviral development efforts may benefit from making note of such calibration issues to produce more accurate models.

**Figure 4:**
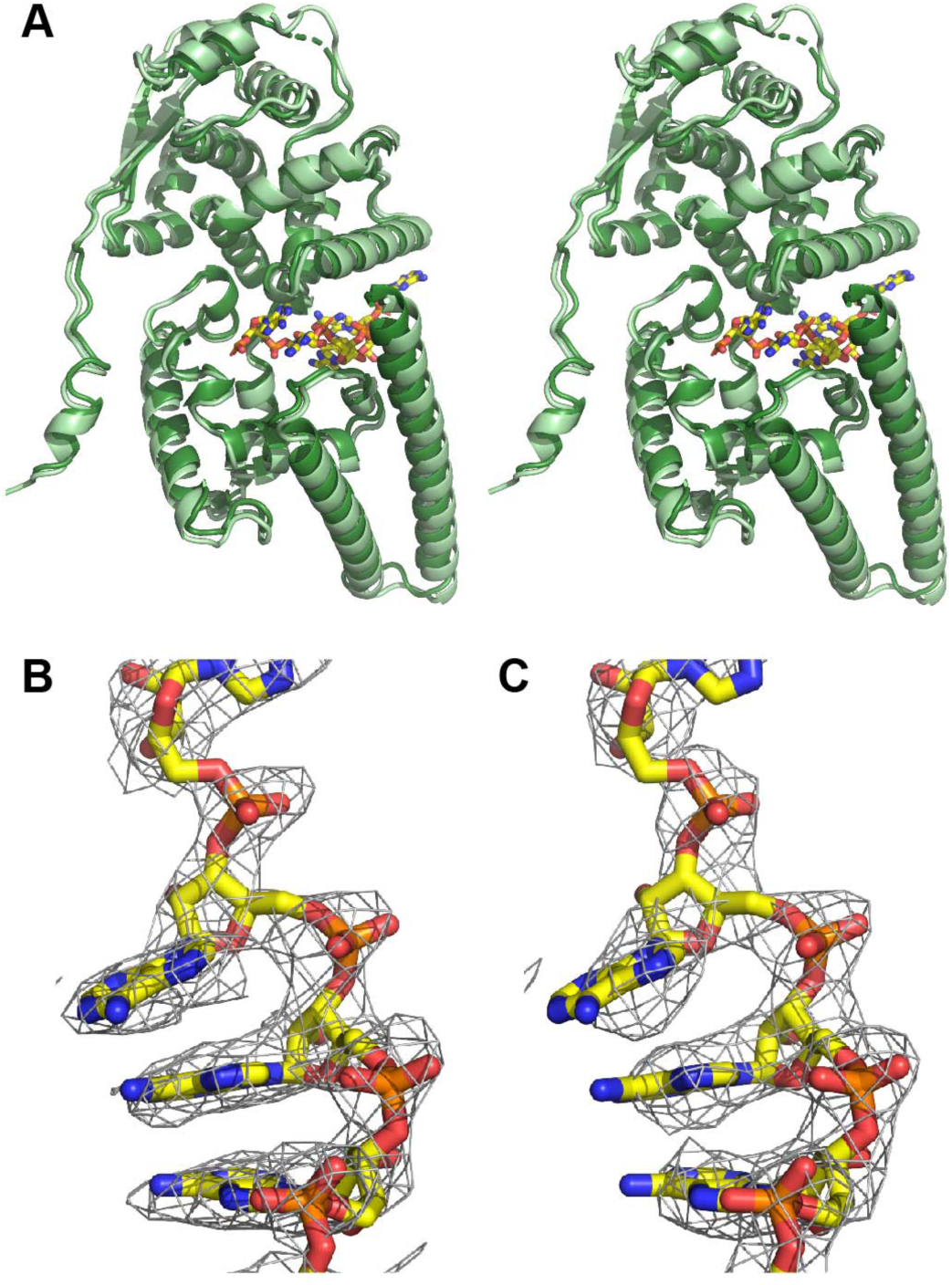
Comparison of NP models. A) Stero view of the 3.1 Å model presented here (dark green) superimposed with the 3.6 Å protein model previously published (light green, 5Z9W.pdb, Sugita *et al.*, 2018). The bound RNA from the model presented here is shown as sticks. Comparison of the 3.1 Å density presented here (B) with that of 3.6 Å density of Sugita *et al.* (2018) (C) shows a clearer definition of the RNA bases in the higher resolution data (B).

Interactions of NP with RNA are primarily mediated by electrostatic interactions where Lys160, Arg174, Arg298 and Arg401 interact directly with negatively charged phosphates of the RNA backbone. Additional residues, Val178, Leu245, Leu331 and Val334, create surfaces for the hydrophobic packing of the nucleotide bases. Oligomerization of NPs is mediated by the N- and C-terminal regions flanking the folded core of the NP (Fig. 3C and D). Ile24 of the NP N-terminal oligomerization region occupies a hydrophobic pocket on an adjacent NP and is expected to directly compete with the VP35 chaperoning peptide for NP binding. The C-terminal oligomerization region folds into a pair of helices which stack on the equivalent helices from adjacent NP using primarily hydrophobic residues. However, turn-on-turn interactions are mediated primarily by charged amino acids which has previously been hypothesized to impart flexibility to the NP filament (Sugita *et al.*, 2018).

## 4. Discussion

The 3.1 Å structure presented here recapitulates the features observed in the previously published 3.6 Å structure (Sugita *et al.*, 2018) including protein-protein and protein-RNA interactions. The improved resolution provides a clear delineation of protein side chains and nucleotide bases clarifying molecular contacts between NP protomers and the RNA. Ebola virus remains an emerging threat to world health security. Key to tackling this threat is the discovery and evaluation of novel therapeutics. The high-resolution structure of the Ebola virus NP bound to RNA will better enable the characterization of potential antiviral drugs.

## Acknowledgements

We gratefully acknowledge Mark Herzik and Bill Anderson for their help in microscope alignment and data collection. A special thanks to Sebastian Raemisch for his help in refining coordinate models using RosettaRelax. We also thank Charles Bowman for computational support.

